# OBE-DB: A Computational Tool and Web Server for the Prediction of Antiobesity Drugs

**DOI:** 10.1101/2025.04.10.648110

**Authors:** Elena Murcia-García, Carlos Martínez-Cortés, Antonio J. Banegas-Luna, Juan José Hernández Morante, Horacio Pérez-Sánchez

## Abstract

The huge obesity prevalence and its associated metabolic disorders highlight the urgent need for new therapeutic strategies beyond lifestyle interventions. Despite the availability of novel pharmacological treatments, the search for more effective and safe anti-obesity compounds remains a challenge. Recent advances in high-performance computational drug discovery have enabled the rapid screening and identification of potential anti-obesity compounds. However, these in silico procedures frequently require complex computational knowledge that limits the use of these techniques for most researchers. To address this gap, we have developed OBE-DB, an accessible and user-friendly platform integrating these computational tools that facilitates the prediction of potential anti-obesity molecules through two complementary approaches: (a) shape similarity analysis against a curated database of approved anti-obesity drugs, and (b) inverse virtual screening of user-submitted molecules against a set of therapeutic protein targets linked to obesity. Our results demonstrate that the server effectively screens and ranks compounds with high predicted activity, outperforming conventional in silico techniques in terms of accuracy and usability. This represents a significant advancement by providing researchers with an intuitive tool to accelerate early-stage drug discovery for obesity treatment. The server is freely accessible without registration, providing users with a detailed report via email upon completion of the predictions. This innovative database and web server is accessible online via http://bio-hpc.eu/software/obe-db/.

## 1. Introduction

Obesity has reached epidemic proportions, with very significant impacts on wellbeing and quality of life and is a major risk factor in other noncommunicable diseases (NCDs). In 2019, it contributed to approximately 5 million deaths from cardiovascular diseases, diabetes, cancers, neurological disorders, chronic respiratory diseases, and digestive disorders (Chong et al., 2023), which constitutes a major public health challenge that also undermines social and economic development throughout the world (Celletti et al., 2024). The fundamental aspect of obesity is the abnormal or excessive body fat accumulation, which is influenced by inherited, physiological, and/or environmental factors, and increases health risk (Bryson et al., 2009).

Several strategies such as calorie restriction, lifestyle management and bariatric surgery have been proposed as therapeutic approaches. Pharmacotherapy has been also employed as a coadjuvant of dietary changes on people with obesity. Some of these drugs include orlistat, which inhibits dietary fat absorption; phentermine-topiramate, and appetite suppressor and satiety enhancer or bupropion-naltrexone, a drug that acts on the brain’s reward system and decreases appetite (Williams et al., 2015). Nonetheless, the development of incretin analogues, like liraglutide, a glucagon-like peptide-1 receptor agonist (GLP-1RA), and semaglutide, a more recent GLP-1 receptor agonist, has changed the game of obesity treatment, showing promising results in clinical trials (Horowitz et al., 2012; Wilding et al., 2021). In fact, other incretins like tirzepatide, a dual GLP-1/GIP receptor agonist has been approved for glycaemic control in type 2 diabetes as well as for obesity management leading in up to 22.5% weight loss in phase 3 obesity trials (Jastreboff et al., 2022). Other combinations of entero-pancreatic hormones including cagrisema (GLP-1/amylin RA) and the triple agonist retatrutide (GLP-1/GIP/glucagon RA) have also progressed to phase 3 trials as obesity treatments and early data suggests that may lead to even greater weight loss than tirzepatide (Melson et al., 2024).

Despite this landscape of recent drugs for obesity treatment, like semaglutide or tirzepatide, there are several limitations that still challenges to effectively addressing this complex condition (Hall & Kahan, 2018). Many antiobesity drugs provide only modest weight loss results and may not be sufficient for those with severe obesity. On the other hand, several side effects that the long-term use of more recent drugs, including gastrointestinal disturbances, increased heart rate, insomnia, and mood changes, and often suicidal ideation has been described with the use of incretin-derived drugs (Tobaiqy & Elkout, 2024; Ueda et al., 2024). Moreover, these drugs are delivered by subcutaneous injection, which is frequently a limitation for several patients. Nevertheless, one of the main caveats of these drugs rely on the fact that individuals may regain weight once they discontinue the medication. Thus, there is a need for new and more effective drugs due to the limitations of existing treatments. This innovation is crucial as the obesity epidemic continues to strain healthcare systems worldwide, necessitating more comprehensive and efficient solutions to improve the overall health and quality of life for individuals struggling with obesity (Melson et al., 2024).

However, incretin-like drugs in isolation will not be enough to address the obesity crisis. Moreover, considering the huge amount of research about obesity molecular mechanisms, it remains a complex and time-consuming task to compile the latest information on potential antiobesity compounds, therapeutic targets and biochemical interactions. Several techniques for drug discovery like ligand similarity experiments and virtual screening for protein-ligand interaction analysis may help in the drug discovery process, saving economic and time resources. Unfortunately, many of this information is evaluated with dedicated bioinformatics tools that are very complex for clinical and experimental researchers. A previously developed web server for drug prediction in diabetes (Pérez-Sánchez et al., 2020) laid the foundation for expanding this approach to anti-obesity compounds, further enhancing predictive capabilities and broadening the scope of the database.

Nevertheless, it is evident that there is a significant gap for a free, easily accessible, and user-friendly online server that serves not only as a database for information on all antiobesity drugs but also as a resource for drug design and development guidance for most researchers. This led us to create the OBE-DB, an innovative online server that, firstly, consolidates and displays comprehensive data on the performance of recognized antiobesity drugs, which allows for ligand-similarity studies. Secondly, OBE-DB serves as a valuable instrument for the virtual docking analyses, essential for the design, advancement, and repurposing of drugs targeting obesity.

## 2. Material and Methods

### 2.1. OBE-DB web server design

The OBE-DB consists of two interconnected tables: Drugs and Calculations. The Drugs table contains data on compounds with potential anti-obesity effects, including their names, chemical formulas, SMILES (Simplified Molecular Input Line Entry System), approval status (approved or experimental), and other relevant information (Table S2). This table serves as the foundation of the database, sharing its core data with the Calculations table.

The Calculations tables store data related to docking experiments and structure similarity comparison requests. These calculations leverage the anti-obesity compound data from the Drugs table. For similarity-based calculations, global three-dimensional shape similarity searches can be conducted using the WEGA (Yan et al., 2013) or SHAFTS (X. Liu et al., 2011) tools. Docking-based calculations are performed using AutoDock Vina (Forli et al., 2016).

OBE-DB is hosted on an Apache 2 server running Ubuntu 18.04.6 64-bit. The user interface is designed to be intuitive and interactive, achieved through the integration of various technologies such as JavaScript (Flanagan, 2011), jQuery (De Volder, 2005), PHP, and HTML. Data managed by OBE-DB is stored in a relational database using the MySQL database management system (Welling & Thomson, 2003). The database is regularly updated, with newly identified compounds being continuously added. Docking and shape similarity queries are processed on a remote computer cluster managed by a SLURM (Yoo et al., 2003) -based job scheduling system. Once completed, results are emailed to the user, including a link to a web page where they are presented in HTML format.

The system is accessible at https://bio-hpc.ucam.edu/obe-db/ without the need for registration. It provides a comprehensive set of resources, including detailed tutorials, examples, frequently asked questions, and related publications.

OBE-DB offers multiple options for querying: searching by compound name or SMILES in the Drugs table or submitting a docking study through the Calculations table. For the former, a detailed view of the resulting compounds is provided online in real time (Figure S1). For docking studies, a detailed report containing the docking prediction results is sent to the user via email once the simulations are completed (Figure S1). This process may take several hours to 2–3 days, depending on the query compound and the workload of the computational cluster. A comprehensive description of the methodology used in designing the Drugs and Calculations tables, as well as a detailed explanation of the experimental workflow, is available in the Supporting Information.

## 3. Results and Discussion

### 3.1. Construction of the database

To investigate the chemical landscape associated with obesity, we developed a database comprising existing drugs for obesity, bioactive compounds with recognized anti-obesity properties, and their corresponding protein targets. Alongside the database, we created tools to facilitate navigation within the anti-obesity chemical space. These tools enable exploration through two distinct approaches: a ligand-based approach and a structure-based approach. In the ligand-based approach, the compound of interest is screened against a library of known anti-obesity drugs.

This ligand-based similarity approach allows for the comparison of the targeted compound— whether a new synthetic molecule, a natural product, or a known drug—with established obesity drugs, potentially revealing novel scaffold similarities that could suggest common functions and activities. To offer flexibility to users, the tool can be accessed either by submitting the SMILES of the desired compound or by drawing the structure of the compound in a dedicated drawing window and submitting it for similarity analysis. The database was manually curated and created to include compounds with known anti-obesity properties.

While the ligand-based similarity approach can suggest potential functions and activities, the structure-based docking approach allows users to assess whether a given compound can interact with the active binding site of a target protein, forming essential interactions (such as hydrogen bonds, van der Waals forces, ionic bonds, and π−π interactions) required for inhibition or activation. This approach may also uncover new amino acids critical for ligand binding, which could enhance the compound’s activity. For the structure-based docking, twenty protein targets associated with obesity were selected (Table S1). These targets can be categorized into four groups based on their role in appetite suppression (e.g., NPYR, MC4R, GLP1R, etc.), regulation of nutrient absorption (e.g., pancreatic lipase, SGLT2), energy expenditure (e.g., ADRA1B, FXR), or inhibition of adipocyte differentiation (e.g., PPARG, FAS, AMPK, etc.). By evaluating the potential effects of compounds on these targets, one can identify the anti-obesity mechanisms of action for a given compound or plant extract. To offer flexibility, the tool can be used by submitting either the SMILES of the desired compound or by drawing its structure in an appropriate window for similarity analysis.

### 3.2. Case studies

To evaluate the functionality of the web server, we conducted similarity and docking calculations using a set of known compounds. These analyses allowed us to assess the accuracy and predictive power of the platform in identifying potential anti-obesity compounds.

#### 3.2.1. Case study 1: Ligand-similarity analyses (Rimonabant)

Rimonabant is an anorectic anti-obesity drug approved by the FDA and EMA in 2006. It is a potent and selective cannabinoid receptor 1 (CB1R) antagonist (Curioni & André, 2006; Pi-Sunyer et al., 2006). Besides its antagonistic properties, numerous studies have shown that, at micromolar concentrations rimonabant behaves as an inverse agonist at CB1 receptors. The drug was the first selective CB1R antagonist/inverse agonist introduced into clinical practice to treat obesity and metabolic-related disorders. It inhibits the proliferation and maturation of adipocytes, improves lipid and glucose metabolism, and regulates food intake and energy balance (Pi-Sunyer et al., 2006). However, it was later withdrawn from market due to CNS-related adverse effects including depression and suicidal ideation (Christensen et al., 2007). Therefore, the search for similar compounds without these adverse effects may be of great interest for drug design (Z. Liu et al., 2021).

To evaluate the functionality of the server, rimonabant structure was introduced on the OBE-DB calculations through the introduction of the SMILES code. After that, a table will be presented containing the sought-after information.

#### 3.2.2. Case study 1: Ligand-similarity analyses (Orforglipron)

Orforglipron has been developed by the pharmaceutical company Lilly and is a next-generation compound belonging to the class of GLP-1 receptor agonists. In 2023, *Lilly* presented the results of phase III clinical trials, demonstrating the effectiveness of this compound in reducing blood glucose levels and promoting weight loss. Although orforglipron has proven effective in treating type 2 diabetes and obesity, it also presents some side effects which are characteristic of GLP-1 agonists, and can induce a delay in gastric emptying, leading to early satiety and gastrointestinal symptoms. Thus, the identification of new molecules could also contribute to a better understanding of the molecular mechanism involved in metabolic control, which is essential for designing more effective and specific therapies for obesity.

To perform the similarity-based calculations, we selected orforglipron as a reference compound. The molecular structure was defined by its canonical SMILES (Simplified Molecular Input Line Entry System) notation, which was entered into the platform to initiate the similarity search. This input served as the basis for identifying structurally related compounds within the database, enabling the prediction of novel molecules with potential anti-obesity activity. Once the calculations were executed, the system returned a ranked list of similar compounds based on their structural resemblance to orforglipron. The results obtained from this analysis are as follows:

The Table 2 presents the results of a ligand-based similarity analysis in OBE-DB, where different compounds were compared to a reference ligand (orforglipron) based on their shape similarity. The columns include Name of the analysed compounds; DB code as a database identifier; Ligand, where structural superposition images of each ligand with the reference compound; and Shape similarity, a numerical value ranging from 0 to 1, where higher values indicate a greater structural resemblance to the reference ligand.

**Table 1.**
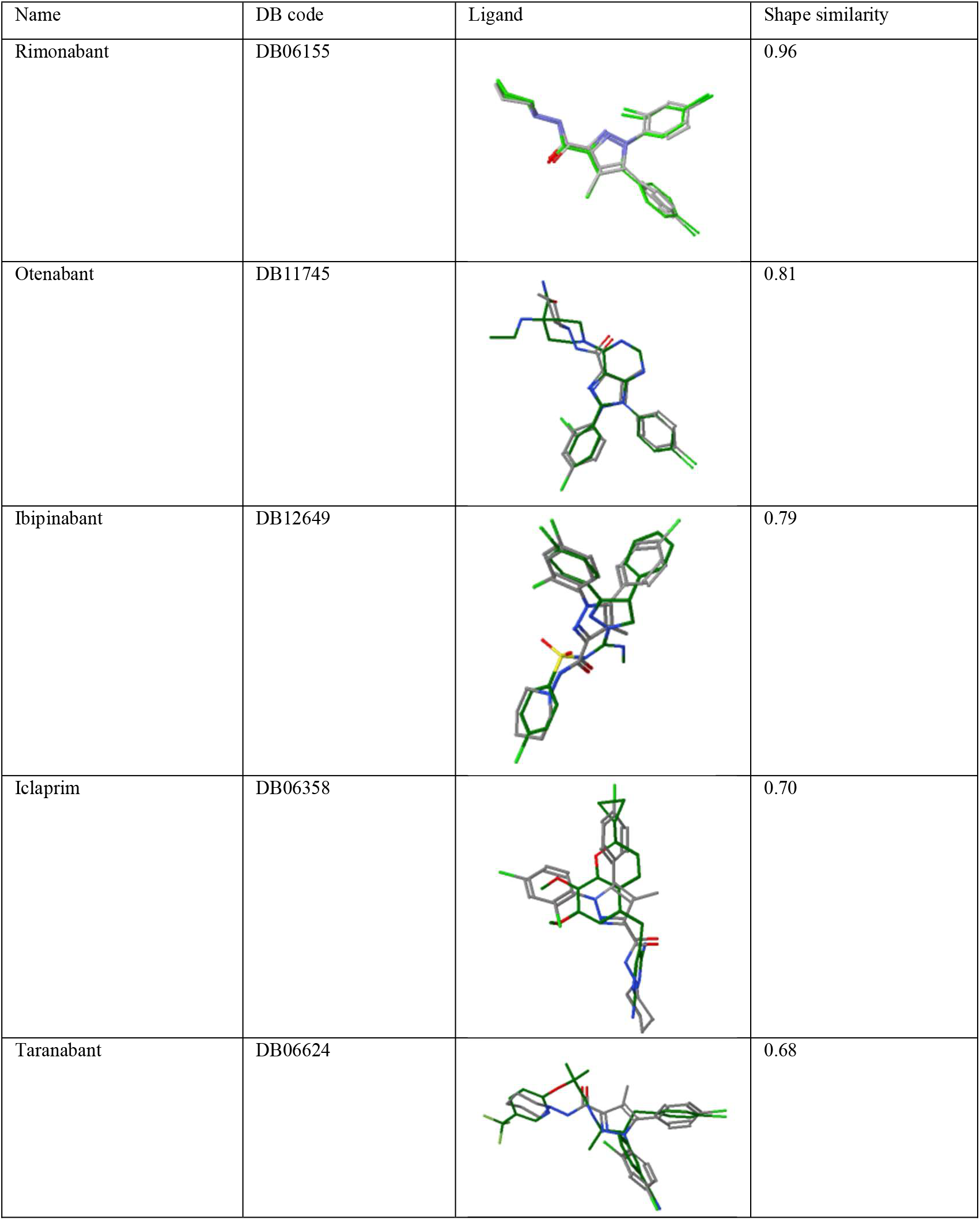
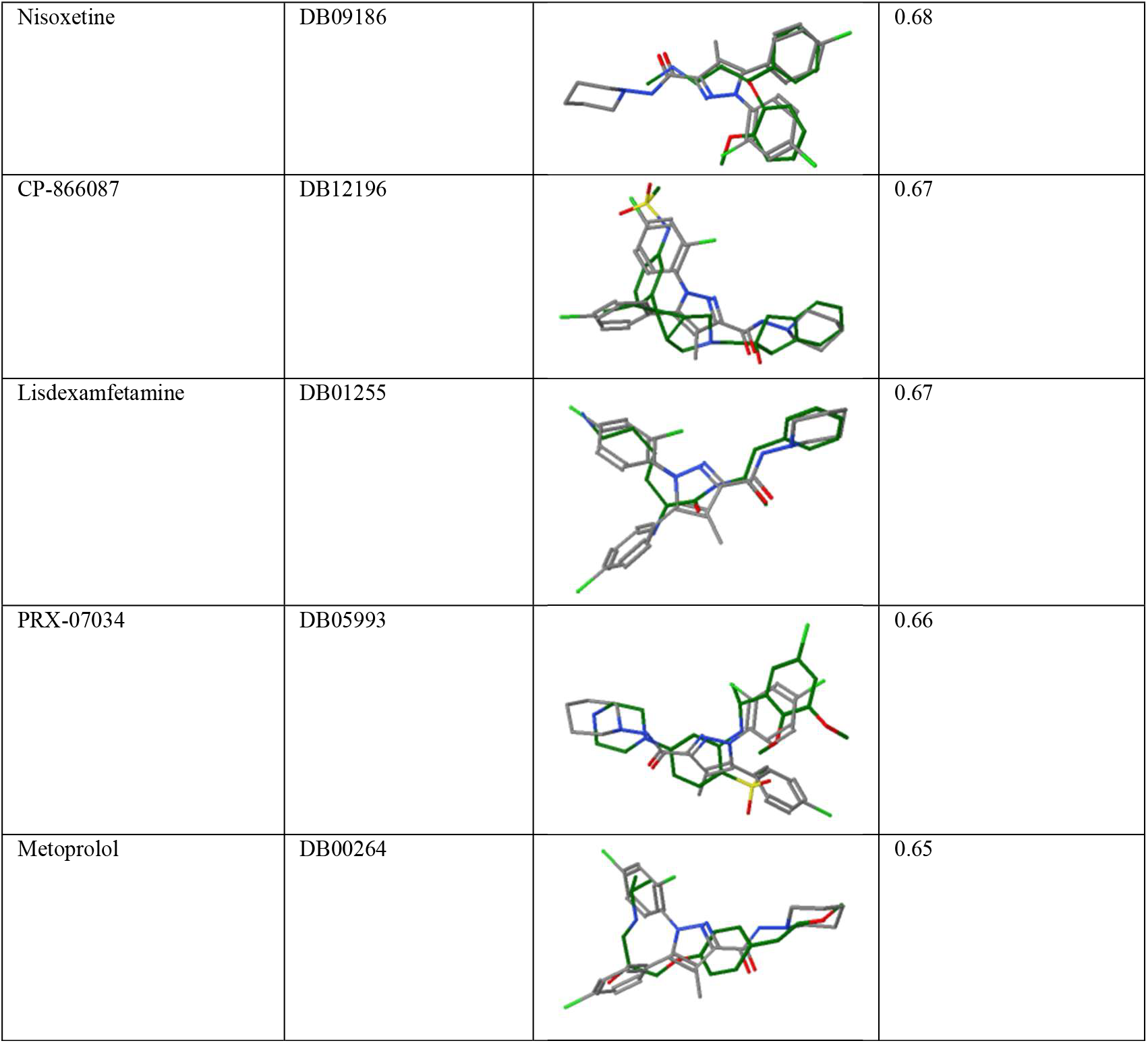
Similarity results for Rimonabant in OBE-DB scored by shape similarity.

| Name | DB code | Ligand | Shape similarity |
| --- | --- | --- | --- |
| Rimonabant | DB06155 |  | 0.96 |
| Otenabant | DB11745 |  | 0.81 |
| Ibipinabant | DB12649 |  | 0.79 |
| Iclaprim | DB06358 |  | 0.70 |
| Taranabant | DB06624 |  | 0.68 |
| Nisoxetine | DB09186 |  | 0.68 |
| CP-866087 | DB12196 |  | 0.67 |
| Lisdexamfetamine | DB01255 |  | 0.67 |
| PRX-07034 | DB05993 |  | 0.66 |
| Metoprolol | DB00264 |  | 0.65 |

**Table 2.**
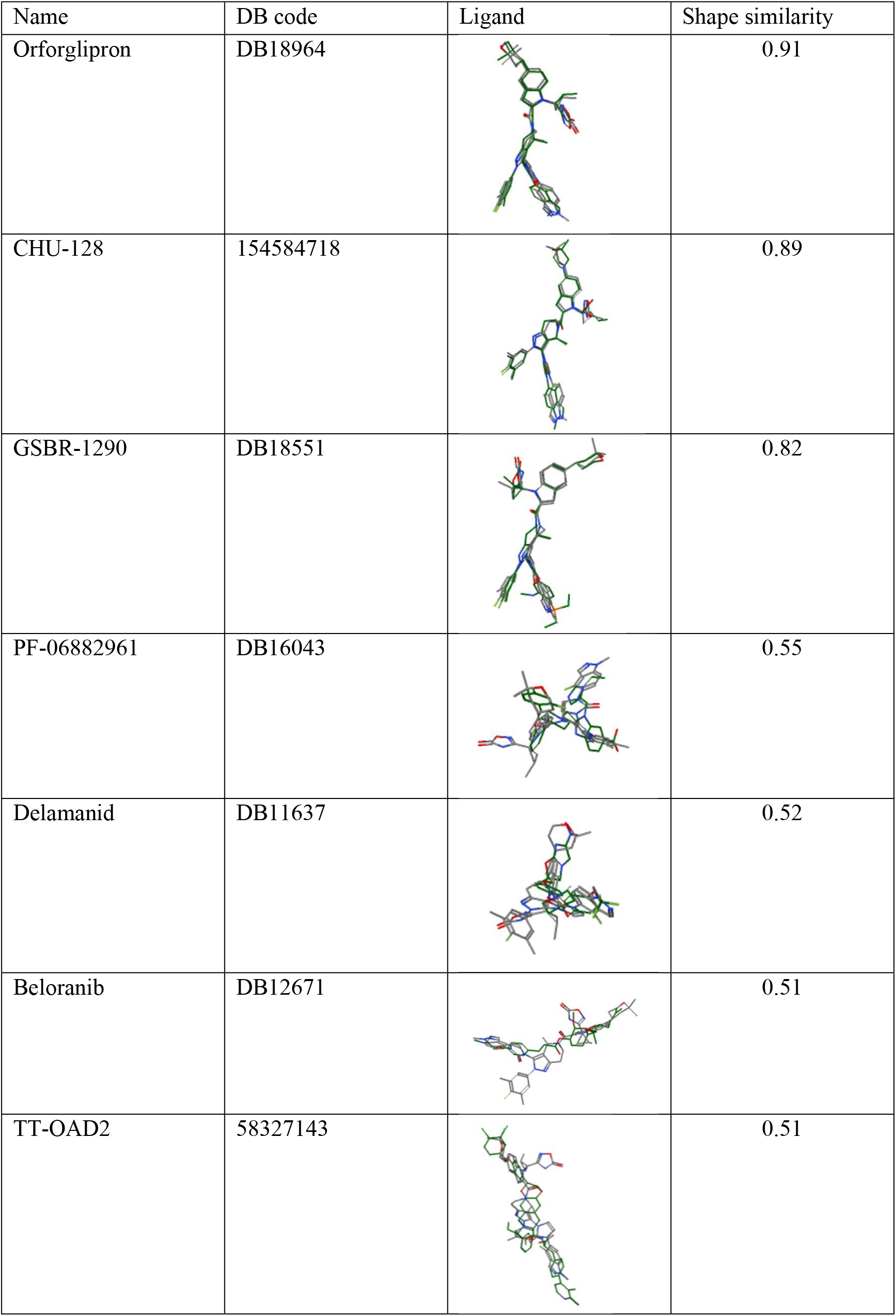
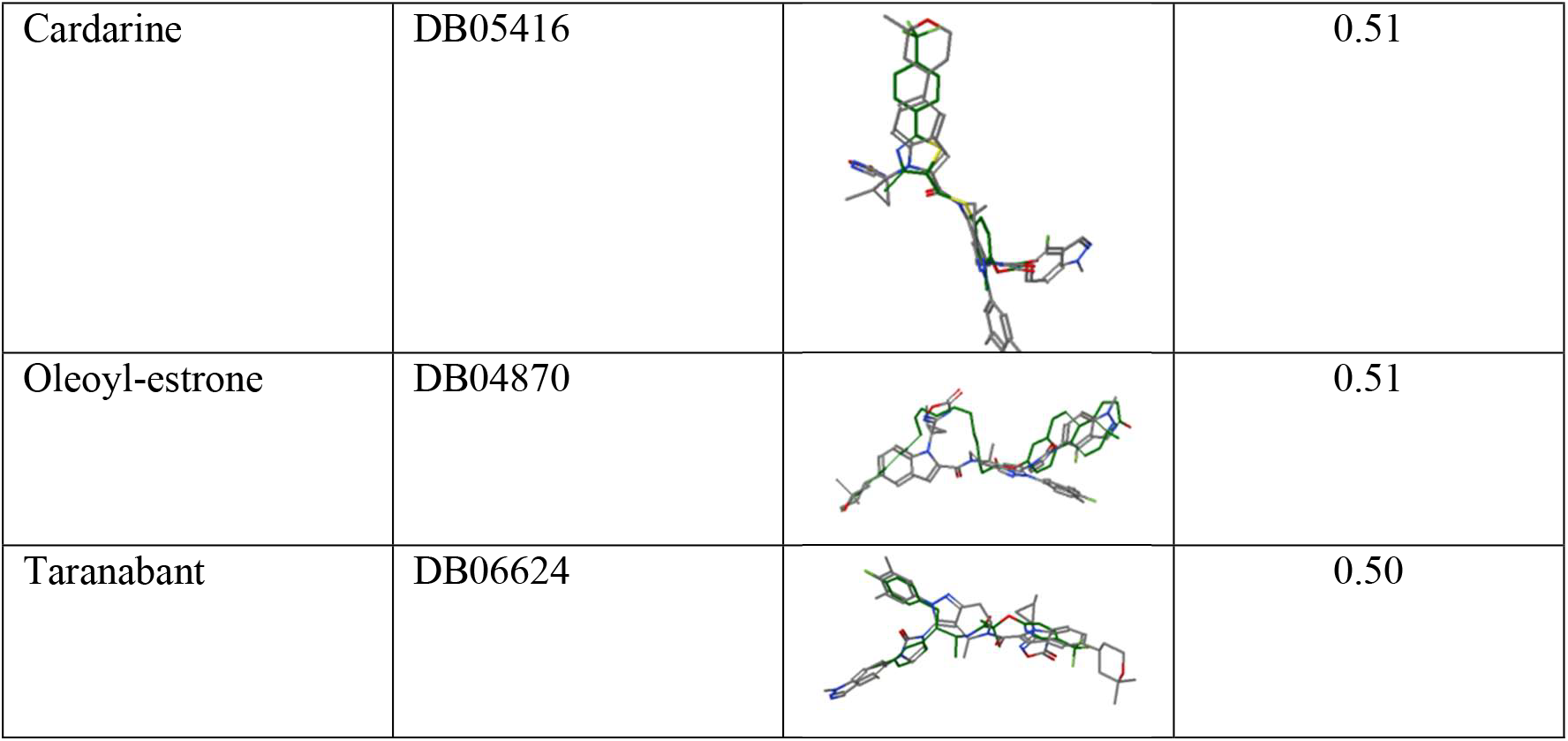
Similarity results for Orforglipron in OBE-DB scored by shape similarity.

| Name | DB code | Ligand | Shape similarity |
| --- | --- | --- | --- |
| Orforglipron | DB18964 |  | 0.91 |
| CHU-128 | 154584718 |  | 0.89 |
| GSBR-1290 | DB18551 |  | 0.82 |
| PF-06882961 | DB16043 |  | 0.55 |
| Delamanid | DB11637 |  | 0.52 |
| Beloranib | DB12671 |  | 0.51 |
| TT-OAD2 | 58327143 |  | 0.51 |
| Cardarine | DB05416 |  | 0.51 |
| Oleoyl-estrone | DB04870 |  | 0.51 |
| Taranabant | DB06624 |  | 0.50 |

The results obtained show that Orforglipron exhibited the highest shape similarity (0.91) with itself, which is an expected result since Orforglipron was used as the query molecule to screen OBE-DB database. Essentially, this means that the method has successfully “found itself”, confirming the accuracy of the similarity-based approach.

CHU-128, other well-defined GLP1R agonist, also showed a high similarity score (0.89), indicating potential structural and functional resemblance, suggesting that three-dimensional structure closely resembles the reference ligand in terms of volume and spatial arrangement. Also, GSBR-1290 displayed a shape similarity of 0.821, implying a moderate structural match with some conformational differences.

#### 3.2.3. Case study 2: Virtual docking analyses (GLP-1R agonist)

Molecular docking calculations were carried out using Orforglipron, a GLP-1R agonist, to evaluate its binding interactions with the receptor. The results revealed that this compound exhibits a high binding affinity, as indicated by a favourable binding energy score, suggesting a strong and stable interaction with the receptor’s active site. Detailed analysis of the docking pose highlights key molecular interactions, including hydrogen bonds and hydrophobic contacts, which may contribute to its potential agonistic activity.

For a more comprehensive understanding of these interactions, users can click on “Additional information” to access a detailed visualization of the protein-ligand complex, including the specific residues involved in binding.

**Figure 1.**
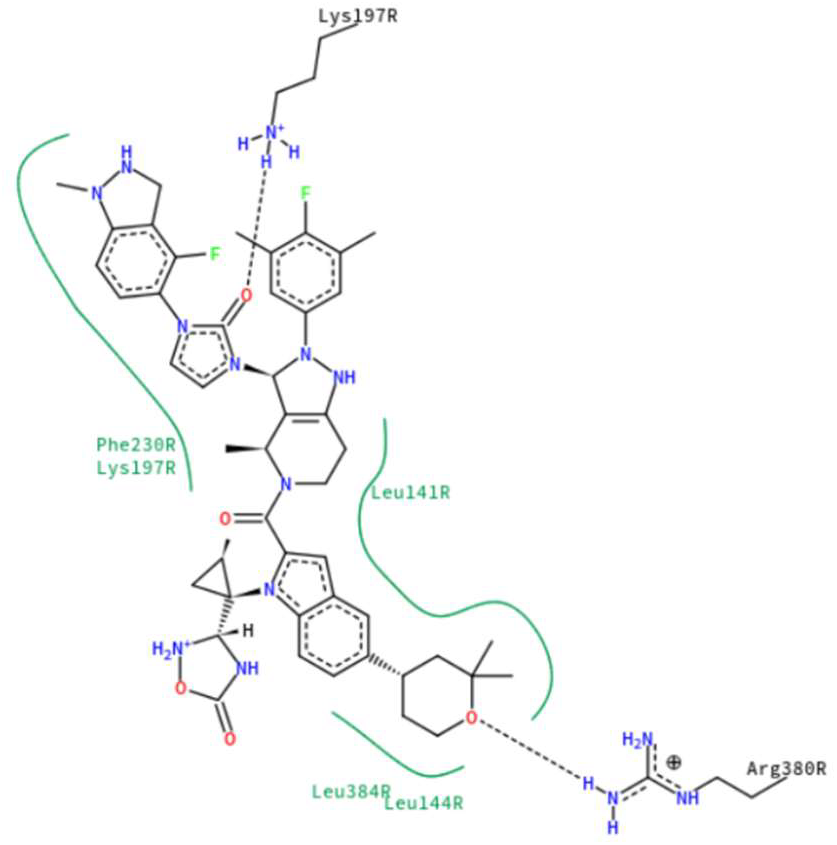
2D schematic representation of the molecular interactions between GLP-1R and the ligand TTOAD2 generated with PoseView. Hydrogen bonds are depicted as dashed lines, and hydrophobic contacts are indicated with green curves. Key interacting residues such as Lys197, Arg380, Leu144, and Phe230 are labelled.

It is also worth noting that PoseView, the external visualization tool integrated into the platform to display ligand-receptor interactions, may occasionally be unable to generate the expected interaction diagram-particularly when the molecular structure or coordinate data are not fully compatible with the rendering requirements. In such cases, the “Additional Information” section becomes especially relevant, as it provides a detailed breakdown of the binding site residues and the nature of their interactions, ensuring that users still have access to critical interpretative data.

#### 3.2.4. Case study 3: Virtual docking analyses (CB1 receptor antagonist)

Rimonabant was also selected as a reference compound to evaluate the predictive performance of OBE-DB web server for molecular docking calculations. The docking results identified CB1R (Cannabinoid Receptor 1) as the top-ranking target, with a binding energy score of –11.2 kcal/mol, suggesting a strong and favourable interaction.

By accessing the “Additional Information” section of the results, it is possible to analyse the specific interactions between Rimonabant and key residues within the CB1R binding pocket. These interactions provide valuable insights into the molecular recognition process and further support the reliability of the docking predictions.

Given these results, Rimonabant proves to be a suitable benchmark compound for testing the functionality of the docking server. Its well-documented activity and strong binding affinity to CB1R make it an appropriate reference for validating the accuracy of computational predictions in the context of anti-obesity drug discovery.

**Figure 2.**
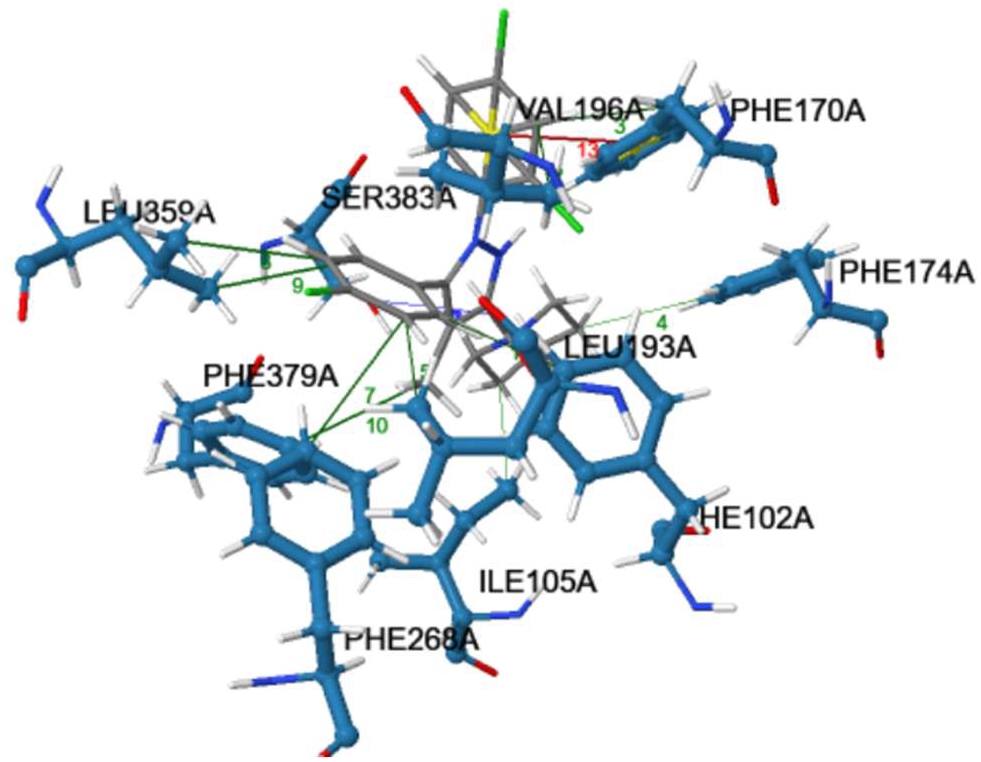
Molecular docking representation of the interactions between the ligand and the active site residues of the target protein (CB1R, PDB: 5TGZ). Hydrophobic interactions are shown in green, *π-π stacking* interactions in red, and hydrogen bonds in blue.

#### 3.2.5. Case study 4: Virtual docking analyses (Pancreatic lipase inhibitor)

In this case, Orlistat was selected as the reference compound to assess the server in predicting pancreatic lipase inhibitors. However, an interesting observation emerged from the docking results, not appearing the pancreatic lipase among the top-ranked targets based on the docking score alone.

This raises an important consideration regarding the limitations of molecular docking and the need to look beyond binding energy predictions when evaluating potential inhibitors. While docking scores serve as a useful first filtering step, they are not always the most reliable indicator of true biological activity. This is particularly relevant for enzymes such as pancreatic lipase, where ligand binding involves not only affinity but also specific interactions with key catalytic residues. Given that Orlistat is a covalent inhibitor, it is likely that the docking algorithm does not fully capture its mechanism of action, leading to an underestimation of its binding affinity.

By accessing the “Additional Information” section of the results, it was possible to analyse the interactions between Orlistat and residues within the active site of pancreatic lipase. Despite its lower docking score, Orlistat still displayed relevant interactions with key catalytic residues, suggesting that it could be a strong inhibitor despite not ranking among the top targets. This highlights the importance of manually reviewing residue-level interactions rather than relying solely on the docking score. Therefore, this study suggests that experimental validation remains crucial for confirming inhibitory activity, particularly for enzymes with complex binding mechanisms. While docking is a valuable technique for preliminary screening, it should always be complemented by residue interaction analysis and biological assays to ensure the identification of truly effective inhibitors.

#### 3.2.6. Case study 5: Virtual docking analyses (PPARγ agonist)

Rosiglitazone is an oral antidiabetic drug belonging to the thiazolidinedione (TZD) class. It functions primarily as a selective agonist of the peroxisome proliferator-activated receptor gamma (PPAR-γ), a nuclear receptor that regulates the expression of genes involved in glucose and lipid metabolism. By activating PPAR-γ, rosiglitazone enhances insulin sensitivity in adipose tissue, skeletal muscle, and the liver, thereby improving glycemic control in patients with type 2 diabetes mellitus (Lehmann et al., 1995; Yki-Järvinen, 2004).

The mechanism of action of rosiglitazone involves modulation of adipocyte differentiation and lipid storage, promoting the uptake and storage of free fatty acids in adipose tissue. This redistribution of lipids from ectopic sites (such as liver and muscle) to adipose tissue contributes to improved insulin sensitivity (Tontonoz & Spiegelman, 2008).

However, the effects of rosiglitazone on body weight and fat distribution have raised concerns. Treatment with rosiglitazone is commonly associated with weight gain, which is thought to result from both increased subcutaneous fat mass and fluid retention (Nissen & Wolski, 2007). While the drug improves insulin resistance, its adipogenic activity—driven by PPAR-γ activation—can promote adipocyte proliferation and lipid accumulation, potentially exacerbating obesity-related outcomes in some patients (Sharma & Staels, 2007).

Due to these effects, rosiglitazone exemplifies the complex interplay between metabolic control and adipogenesis, highlighting the need for therapeutic strategies that enhance insulin sensitivity without promoting excessive weight gain.

We selected rosiglitazone to further assess the performance of the docking module implemented in the web server. This molecule is a well-known ligand with reported affinity for the peroxisome proliferator-activated receptor gamma (PPAR-γ). However, in this case, PPAR-γ was not ranked among the top predicted targets. This outcome highlights a known limitation of molecular docking approaches: while generally effective, they do not guarantee accurate predictions in every scenario. The current implementation relies on the standard version of Autodock Vina, which has demonstrated robust performance across a wide range of targets but is not universally reliable. As reported in several benchmarking studies and review articles, docking algorithms, including Vina, may occasionally misrank targets due to factors such as protein flexibility, scoring function limitations, or suboptimal ligand orientation (Trott & Olson, 2010).

**Figure 3.**
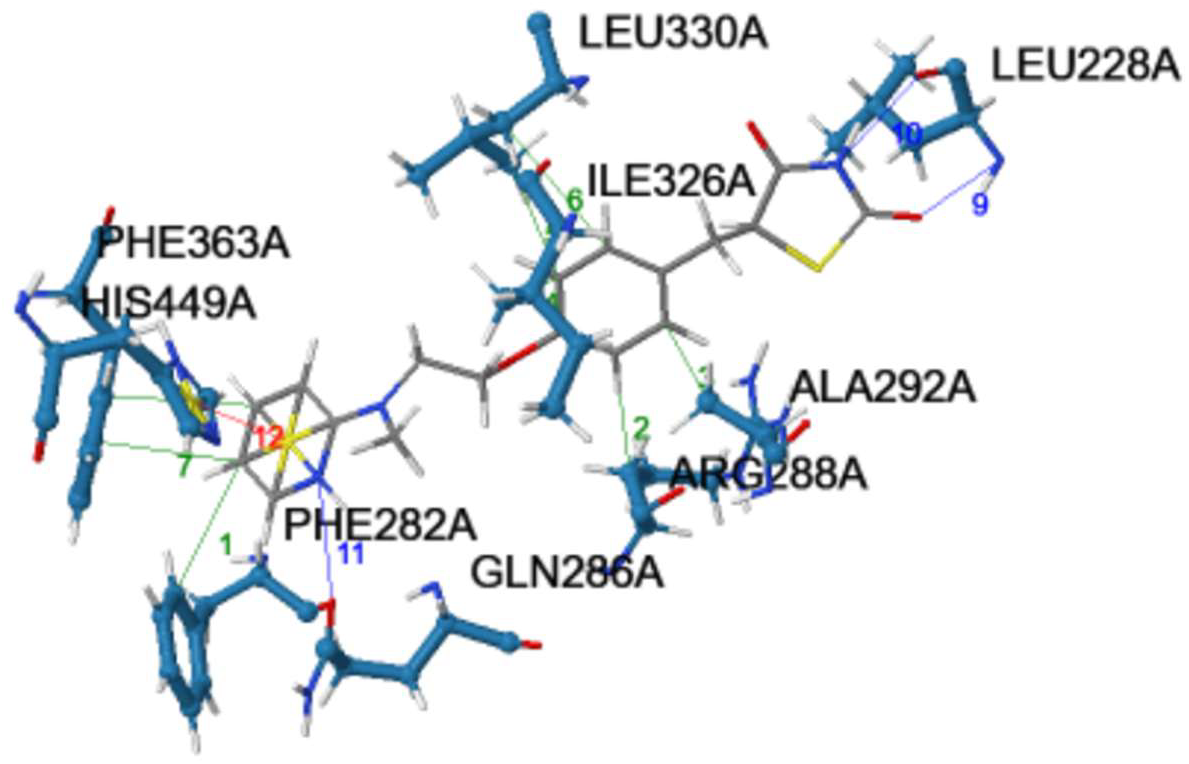
Molecular docking representation of the interactions between the ligand and the active site residues of the target protein (PPAR-γ receptor, PDB: 2FVJ). Hydrophobic interactions are shown in green, *π-π stacking* interactions in red, and hydrogen bonds in blue.

## 4. Conclusion

In this study, we present the development of a novel chemoinformatic server, OBE-DB, which consolidates data on drugs and proteins involved in obesity. Using this information, we have developed a variety of tools that users can employ to aid in the design of anti-obesity drugs. This platform allows users to screen compounds using either a ligand-based approach, based on molecular similarity analysis, or a structure-based approach, leveraging molecular docking simulations. By providing these complementary methodologies, the server enhances flexibility in identifying bioactive molecules with potential therapeutic applications. To assess the functionality of the server, we conducted similarity calculations and docking simulations using representative compounds targeting different anti-obesity mechanisms. Our results demonstrate that the platform effectively identifies structurally and functionally relevant molecules, providing reliable predictions across multiple therapeutic targets. The docking analyses confirmed the ability of the server to prioritize compounds with high binding affinity, aligning with established pharmacological profiles. We envision that our server will play a key role in navigating the chemical space towards the discovery of new anti-obesity drugs or the repurposing of existing ones.

The integration of chemoinformatic and computational modelling into a user-friendly interface enhances the accessibility of advanced in silico screening tools for the scientific community. By bridging the gap between computational predictions and experimental validation, this platform streamlines the identification of promising anti-obesity candidates and supports early-stage drug discovery.

Future developments will focus on expanding the compound database refining the predictive algorithms and incorporating additional obesity-related targets to further improve the accuracy and applicability of the tool. Ultimately, this web server serves as a valuable resource for researchers in the field of obesity drug discovery, facilitating the rapid screening of bioactive compounds and accelerating the development of novel therapeutic strategies.

## 5. CRediT authorship contribution statement

**Elena Murcia-García:** Conceptualization, Formal analysis, Investigation, Validation, Writing – original draft, Writing – review & editing. **Carlos Martínez-Cortés**: Conceptualization, Data curation, Formal analysis, Methodology, Software, Validation, Writing – review & editing. **Antonio Jesús Banegas-Luna**: Conceptualization, Data curation, Methodology, Software, Writing – review & editing. ***Juan José Hernández-Morante***: Conceptualization, Investigation, Supervision, Validation, Writing – review & editing. ***Horacio Pérez-Sánchez***: Conceptualization, Funding acquisition, Methodology, Project administration, Resources, Supervision, Writing – review & editing.

## 7. Funding and Acknowledgments

This research was partially supported by the supercomputing infrastructure of the NLHPC (CCSS210001).

## Supplemental

The supporting information is available under request to corresponding authors. Further details on methodology implemented in OBE-DB database design; Table S1:

Pharmaceutical targets of obesity for SBVS in the OBE-DB Experiments database; Table S2:

Number of drugs curated in the OBE-DB Drugs; Supporting Figures S1: OBE-DB web server user interface; Supporting data Table S2 for anti-obesity compounds of the Drugs table (XLSX)

## References

Bryson, A, De La Motte, S., & Dunk, C. (2009). Reduction of dietary fat absorption by the novel gastrointestinal lipase inhibitor cetilistat in healthy volunteers. British Journal of Clinical Pharmacology, 67(3), 309–315. 10.1111/J.1365-2125.2008.03311.X

Celletti, F., Branca, F., & Farrar, J. (2024). Obesity and Glucagon-Like Peptide-1 Receptor Agonists. JAMA. 10.1001/JAMA.2024.25872

Chong, B., Jayabaskaran, J., Kong, G., Chan, Y. H., Chin, Y. H., Goh, R., Kannan, S., Ng, C. H., Loong, S., Kueh, M. T. W., Lin, C., Anand, V. V., Lee, E. C. Z., Chew, H. S. J., Tan, D. J. H., Chan, K. E., Wang, J. W., Muthiah, M., Dimitriadis, G. K., … Chew, N. W. S. (2023). Trends and predictions of malnutrition and obesity in 204 countries and territories: an analysis of the Global Burden of Disease Study 2019. EClinicalMedicine, 57, 101850. 10.1016/j.eclinm.2023.101850

Christensen, R., Kristensen, P. K., Bartels, E. M., Bliddal, H., & Astrup, A. (2007). Efficacy and safety of the weight-loss drug rimonabant: a meta-analysis of randomised trials. Lancet (London, England), 370(9600), 1706–1713. 10.1016/S0140-6736(07)61721-8

Curioni, C., & André, C. (2006). Rimonabant for overweight or obesity. The Cochrane Database of Systematic Reviews, 2006(4). 10.1002/14651858.CD006162.PUB2

De Volder, K. (2005). JQuery: A generic code browser with a declarative configuration language. Lecture Notes in Computer Science (Including Subseries Lecture Notes in Artificial Intelligence and Lecture Notes in Bioinformatics), 3819 LNCS, 88–102. 10.1007/11603023_7

Flanagan, D. (2011). JavaScript: The Definitive Guide 6th Edition. Chemistry, 994. https://books.google.com/books/about/JavaScript_The_Definitive_Guide.html?hl=es&id=4RChxt67lvwC

Forli, S., Huey, R., Pique, M. E., Sanner, M. F., Goodsell, D. S., & Olson, A. J. (2016). Computational protein-ligand docking and virtual drug screening with the AutoDock suite. Nature Protocols, 11(5), 905–919. 10.1038/NPROT.2016.051

Hall, K. D., & Kahan, S. (2018). Maintenance of lost weight and long-term management of obesity. The Medical Clinics of North America, 102(1), 183. 10.1016/J.MCNA.2017.08.012

Horowitz, M., Flint, A., Jones, K. L., Hindsberger, C., Rasmussen, M. F., Kapitza, C., Doran, S., Jax, T., Zdravkovic, M., & Chapman, I. M. (2012). Effect of the once-daily human GLP-1 analogue liraglutide on appetite, energy intake, energy expenditure and gastric emptying in type 2 diabetes. Diabetes Research and Clinical Practice, 97(2), 258–266. 10.1016/J.DIABRES.2012.02.016

Jastreboff, A. M., Aronne, L. J., Ahmad, N. N., Wharton, S., Connery, L., Alves, B., Kiyosue, A., Zhang, S., Liu, B., Bunck, M. C., & Stefanski, A. (2022). Tirzepatide Once Weekly for the Treatment of Obesity. The New England Journal of Medicine, 387(3), 205–216. 10.1056/NEJMOA2206038

Lehmann, J. M., Moore, L. B., Smith-Oliver, T. A., Wilkison, W. O., Willson, T. M., & Kliewer, S. A. (1995). An antidiabetic thiazolidinedione is a high affinity ligand for peroxisome proliferator-activated receptor gamma (PPAR gamma). The Journal of Biological Chemistry, 270(22), 12953–12956. 10.1074/JBC.270.22.12953

Liu, X., Jiang, H., & Li, H. (2011). SHAFTS: A hybrid approach for 3D molecular similarity calculation. 1. method and assessment of virtual screening. Journal of Chemical Information and Modeling, 51(9), 2372–2385. 10.1021/CI200060S/SUPPL_FILE/CI200060S_SI_001.PDF

Liu, Z., Iyer, M. R., Godlewski, G., Jourdan, T., Liu, J., Coffey, N. J., Zawatsky, C. N., Puhl, H. L., Wess, J., Meister, J., Liow, J. S., Innis, R. B., Hassan, S. A., Lee, Y. S., Kunos, G., & Cinar, R. (2021). Functional Selectivity of a Biased Cannabinoid-1 Receptor (CB1R) Antagonist. ACS Pharmacology and Translational Science, 4(3), 1175–1187. 10.1021/ACSPTSCI.1C00048/SUPPL_FILE/PT1C00048_SI_001.PDF

Melson, E., Ashraf, U., Papamargaritis, D., & Davies, M. J. (2024). What is the pipeline for future medications for obesity? International Journal of Obesity 2024, 1–19. 10.1038/s41366-024-01473-y

Pérez-Sánchez, H., den-Haan, H., Peña-García, J., Lozano-Sánchez, J., Martínez Moreno, M.E., Sánchez-Pérez, A., Muñoz, A., Ruiz-Espinosa, P., Pereira, A. S. P., Katsikoudi, A., Gabaldón Hernández, J. A., Stojanovic, I., Carretero, A. S., & Tzakos, A. G. (2020). DIA-DB: A Database and Web Server for the Prediction of Diabetes Drugs. Journal of Chemical Information and Modeling, 60(9), 4124–4130. 10.1021/ACS.JCIM.0C00107

Pi-Sunyer, F. X., Aronne, L. J., Heshmati, H. M., Devin, J., & Rosenstock, J. (2006). Effect of rimonabant, a cannabinoid-1 receptor blocker, on weight and cardiometabolic risk factors in overweight or obese patients - RIO-North America: A randomized controlled trial. JAMA, 295(7), 761–775. 10.1001/jama.295.7.761

Sharma, A. M., & Staels, B. (2007). Review: Peroxisome proliferator-activated receptor gamma and adipose tissue--understanding obesity-related changes in regulation of lipid and glucose metabolism. The Journal of Clinical Endocrinology and Metabolism, 92(2), 386–395. 10.1210/JC.2006-1268

Tobaiqy, M., & Elkout, H. (2024). Psychiatric adverse events associated with semaglutide, liraglutide and tirzepatide: a pharmacovigilance analysis of individual case safety reports submitted to the EudraVigilance database. International Journal of Clinical Pharmacy, 46(2), 488. 10.1007/S11096-023-01694-7

Tontonoz, P., & Spiegelman, B. M. (2008). Fat and beyond: the diverse biology of PPARgamma. Annual Review of Biochemistry, 77, 289–312. 10.1146/ANNUREV.BIOCHEM.77.061307.091829

Trott, O., & Olson, A. J. (2010). AutoDock Vina: improving the speed and accuracy of docking with a new scoring function, efficient optimization and multithreading. Journal of Computational Chemistry, 31(2), 455. 10.1002/JCC.21334

Ueda, P., Söderling, J., Wintzell, V., Svanström, H., Pazzagli, L., Eliasson, B., Melbye, M., Hviid, A., & Pasternak, B. (2024). GLP-1 Receptor Agonist Use and Risk of Suicide Death. JAMA Internal Medicine, 184(11), 1301–1312. 10.1001/JAMAINTERNMED.2024.4369

Update on FDA’s ongoing evaluation of reports of suicidal thoughts or actions in patients taking a certain type of medicines approved for type 2 diabetes and obesity | FDA. (n.d.). Retrieved December 17, 2024, from https://www.fda.gov/drugs/drug-safety-and-availability/update-fdas-ongoing-evaluation-reports-suicidal-thoughts-or-actions-patients-taking-certain-type?utm_source=chatgpt.com

Welling, Luke., & Thomson, Laura. (2003). PHP and MySQL Web development. 871.

Wilding, J. P. H., Batterham, R. L., Calanna, S., Davies, M., Van Gaal, L. F., Lingvay, I., McGowan, B. M., Rosenstock, J., Tran, M. T. D., Wadden, T. A., Wharton, S., Yokote, K., Zeuthen, N., & Kushner, R. F. (2021). Once-Weekly Semaglutide in Adults with Overweight or Obesity. The New England Journal of Medicine, 384(11), 989–1002. 10.1056/NEJMOA2032183

Williams, E. P., Mesidor, M., Winters, K., Dubbert, P. M., & Wyatt, S. B. (2015). Overweight and Obesity: Prevalence, Consequences, and Causes of a Growing Public Health Problem. Current Obesity Reports, 4(3), 363–370. 10.1007/S13679-015-0169-4/METRICS

Yan, X., Li, J., Liu, Z., Zheng, M., Ge, H., & Xu, J. (2013). Enhancing molecular shape comparison by weighted Gaussian functions. Journal of Chemical Information and Modeling, 53(8), 1967–1978. 10.1021/CI300601Q/SUPPL_FILE/CI300601Q_SI_001.PDF

Yki-Järvinen, H. (2004). Thiazolidinediones. New England Journal of Medicine, 351(11), 1106– 1118. 10.1056/NEJMRA041001

Yoo, A. B., Jette, M. A., & Grondona, M. (2003). SLURM: Simple Linux Utility for Resource Management. Lecture Notes in Computer Science (Including Subseries Lecture Notes in Artificial Intelligence and Lecture Notes in Bioinformatics), 2862, 44–60. 10.1007/10968987_3

